# Uncovering the link between ATP synthase and the TCA cycle by crosslinking mass spectrometry

**DOI:** 10.64898/2025.12.13.694125

**Authors:** Laura Pérez Pañeda, Jelena Misic, Tereza Kadavá, Nils-Göran Larsson, Albert J.R. Heck

## Abstract

Mitochondrial oxidative phosphorylation (OXPHOS) comprises a series of multi-subunit protein complexes that operate in coordination with the tricarboxylic acid (TCA) cycle to generate ATP. Although these systems are metabolically interconnected, complex II is generally regarded as the only direct structural link between the OXPHOS and TCA cycle. Here, we combine in-solution crosslinking mass-spectrometry (XL-MS), quantitative proteomics, and blue native PAGE (BN-PAGE) to explore how ATP synthase (complex V) integrates within the mitochondrial metabolic network under physiological and pathological conditions. We demonstrate that in murine wild-type hearts, the F_1_ catalytic head of ATP synthase forms extensive contacts with TCA cycle enzymes, establishing a previously unanticipated link between the OXPHOS and central carbon metabolism. We also found that under mitochondrial dysfunction, in this case LRPPRC-deficient hearts, where defective mitochondrial DNA gene expression destabilizes ATP synthase, these interactions become strengthened. Moreover, ATP synthase dysfunction promotes binding of the ATPase inhibitory factor 1 (ATIF1) to the F_1_ head via its N-terminal inhibitory region, shifting the ATP synthase toward an energy-preserving state. Together, our findings show that ATP synthase deficiency drives remodeling of the F_1_ interactome, revealing how mitochondrial structure and regulation adapt to preserve energy homeostasis under stress.

## Introduction

Mitochondria are essential organelles present in nearly all eukaryotic cells, involved in energy conversion, metabolism, and signaling. Their main bioenergetic function is carried out by the oxidative phosphorylation (OXPHOS) system, which generates the majority of cellular ATP. OXPHOS comprises the respiratory chain complexes (I–IV) and ATP synthase (complex V), all embedded in the inner mitochondrial membrane. In mammals, it is well established that respiratory chain complexes can assemble into higher-order supramolecular structures known as supercomplexes (Milenkovic *et al*, 2017; Fernández-Vizarra & Ugalde, 2022). However, the importance of these assemblies for mitochondrial bioenergetics has been challenged (Milenkovic *et al*, 2023; Fernández-Vizarra *et al*, 2022) and their *in vivo* role remains to be fully elucidated. By contrast, ATP synthase forms higher-order oligomers that play a well-defined architectural role in shaping the inner mitochondrial membrane (Blum *et al*, 2019; Kühlbrandt, 2019). The ATP synthase monomers associate to form dimers, which further arrange into rows along the tips of cristae membranes, promoting inner membrane curvature shaping cristae morphology.

In addition to its architectural role, ATP synthase serves as a central energy-conversion machinery in mitochondria. It uses the proton-motive force generated by the respiratory chain (complexes I, III and IV) to synthesize ATP through a rotary catalytic mechanism (Abrahams *et al*, 1994; Boyer, 1993; Noji *et al*, 1997). Each monomer is composed of two major domains, the soluble F_1_ catalytic head and the membrane-embedded F_O_ proton channel, which are connected by the peripheral stalk (Walker, 2013; Spikes *et al*, 2020). When the proton-motive force collapses due to mitochondrial depolarization or respiratory chain dysfunction, ATP synthase can reverse its direction and hydrolyze ATP to help maintain the inner membrane potential (Buchet & Godinot, 1998). This reverse activity is tightly regulated by the endogenous inhibitory factor ATIF1, a small protein (∼10 kDa), which binds the F_1_ head in a pH-dependent manner and prevents wasteful ATP hydrolysis (Cabezon *et al*, 2000; Gledhill *et al*, 2007).

The function of ATP synthase rely on subunits and assembly factors encoded by two genetic systems, namely the nuclear and mitochondrial genomes (mtDNA). Because of this dual genetic origin, the synthesis and import of nuclear-encoded components must be coordinated with the expression of mtDNA-encoded subunits to ensure accurate assembly and maturation of the complex. Mutations in ATP synthase subunits, defects affecting mtDNA maintenance or gene expression, and mutations in assembly or regulatory factors can all compromise the integrity of the ATP synthase complex, leading to the accumulation of incomplete intermediates (Mourier *et al*, 2014; Milenkovic *et al*, 2013; Čížková *et al*, 2008; Vrbacký *et al*, 2016; Silva-Pinheiro *et al*, 2023). These molecular defects ultimately impair ATP synthase activity, cause mitochondrial dysfunction, and contribute to the pathogenesis of numerous human disorders, including cardiovascular, neurodegenerative, and neurodevelopmental diseases (Galber *et al*, 2021).

ATP synthase operates in tight bioenergetic cooperation with both the respiratory chain complexes and the tricarboxylic acid (TCA) cycle, which provides reducing equivalents to sustain electron flow and proton-motive force generation. Complex II (succinate dehydrogenase) is unique in this regard, as it functions simultaneously as a TCA cycle enzyme and as an integral component of the OXPHOS system, thereby forming a direct structural and functional bridge between the two pathways. Electrons from NADH, produced by multiple oxidative steps of the TCA cycle, are transferred to the respiratory chain through complex I, whereas those from succinate oxidation enter via complex II. This metabolic coupling is essential for efficient energy conversion; however, it has remained elusive whether additional physical connections, besides those involving complex II, exist between the TCA cycle enzymes and the OXPHOS system.

For long, it has been hypothesized that metabolic enzymes, including those of the TCA cycle, can associate to form metabolons to facilitate substrate channeling and coordinate metabolic flux through interconnected pathways (Elcock & McCammon, 1996; Morgunov & Srere, 1998; Srere, 1985). However, such interactions are typically transient, dynamic, and weak, preventing their detection by most conventional biochemical or structural approaches. To capture these interactions and determine their spatial organization, advanced structural proteomics approaches are required. In-solution crosslinking mass spectrometry (XL-MS) has recently emerged as a powerful tool to investigate protein organization within intact mitochondria (Liu *et al*, 2018; Hevler & Heck, 2023; Schuhmann *et al*, 2025; Schweppe *et al*, 2017; Linden *et al*, 2020; Caudal *et al*, 2022). By covalently linking proximal amino acid residues, this technique enables direct mapping of protein-protein interactions arising from close spatial proximity. While most biochemical and structural methods require detergents and partial mitochondrial solubilization, XL-MS preserves the native environment in mitochondria, capturing both stable and transient interactions at a proteome-wide scale (Piersimoni *et al*, 2022; Wheat *et al*, 2021). This approach therefore provides a unique opportunity to uncover functional protein networks *in situ*, within their physiological context.

Here, we applied in-solution XL-MS to investigate the (re)organization of the OXPHOS interaction network in the mouse heart under different conditions. Although extensive structural information exists for individual OXPHOS complexes and their higher-order assemblies (Hevler & Heck, 2023; Boekema & Braun, 2007; Zheng *et al*, 2024), much less is known about how these complexes interact with other metabolic pathways within mitochondria. Performing XL-MS on intact heart mitochondria, we mapped protein interactions under normal physiological conditions and in a conditional *Lrpprc* knockout mouse model with impaired mtDNA gene expression in the heart. Our XL-MS analyses revealed previously unrecognized associations between the F_1_ domain of ATP synthase and multiple TCA cycle enzymes in wild-type hearts. These interactions were further enhanced under mitochondrial dysfunction caused by compromised ATP synthase integrity in the *Lrpprc* knockout hearts. In addition to the TCA cycle enzymes, several other metabolic enzymes also showed increased association with the F_1_ head in the absence of LRPPRC. Notably, *in vivo* binding of the N-terminal inhibitory domain of ATIF1 to the F_1_ head of ATP synthase was observed exclusively in the *Lrpprc* knockouts. Our findings provide a link between the structural remodeling of ATP synthase and the inhibitory activation of ATIF1 under mitochondrial stress conditions.

## Results

### Mitochondrial ATP synthase associates with the TCA cycle enzymes in the mouse heart

To gain a broader understanding of the spatial organization of OXPHOS in mitochondria, we first revisited published in-solution XL-MS data from intact murine heart mitochondria (Milenkovic *et al*, 2023). While the original analyses primarily focused on intra- and inter-complex interactions within the respirasome (the respiratory chain supercomplex I-III_2_-IV) in wild-type and respirasome-deficient hearts, the underlying dataset contained valuable information extending beyond the respiratory chain, providing an opportunity to explore additional mitochondrial protein associations.

Here, we focused especially on identified crosslinks between OXPHOS complexes and central metabolic pathways within mitochondria. The wild-type mouse heart mitochondria dataset revealed extensive interactions between the OXPHOS complexes and enzymes of the TCA cycle (**Figures 1A and S1**). Although complex II has been considered the primary structural bridge between these two systems, the ATP synthase exhibited the strongest connectivity to TCA cycle enzymes among all OXPHOS complexes (**Figures 1A and S1)**. Crosslinks were detected across both the soluble F_1_ and membrane-embedded F_O_ subunits of ATP synthase, and involved multiple enzymes of the TCA cycle, including most notably citrate synthase (CISY), isocitrate dehydrogenase isoforms (IDHP, IDH3A), 2-oxoglutarate dehydrogenase subunits (ODPA, ODO1, ODO2), succinyl-CoA ligase subunits (SUCA, SUCB1), fumarate hydratase (FUMH) and malate dehydrogenase (MDHM) (**Figure 1A and S1**). Most detected crosslinks of TCA cycle enzymes to ATP synthase were formed with subunits of the F_1_ catalytic head, primarily the α (ATPA) and β (ATPB) subunits (**Figures 1A and S1**). Interestingly, the number of such crosslinks was roughly twofold lower than those within the ATP synthase itself, yet about twice as abundant as the crosslinks detected between individual TCA cycle enzymes (**Figure S1**). Collectively, these data suggest that OXPHOS and the TCA cycle are structurally and functionally closely coupled in cardiac mitochondria, with ATP synthase positioned at the key interface between these two systems.

**Figure 1.**
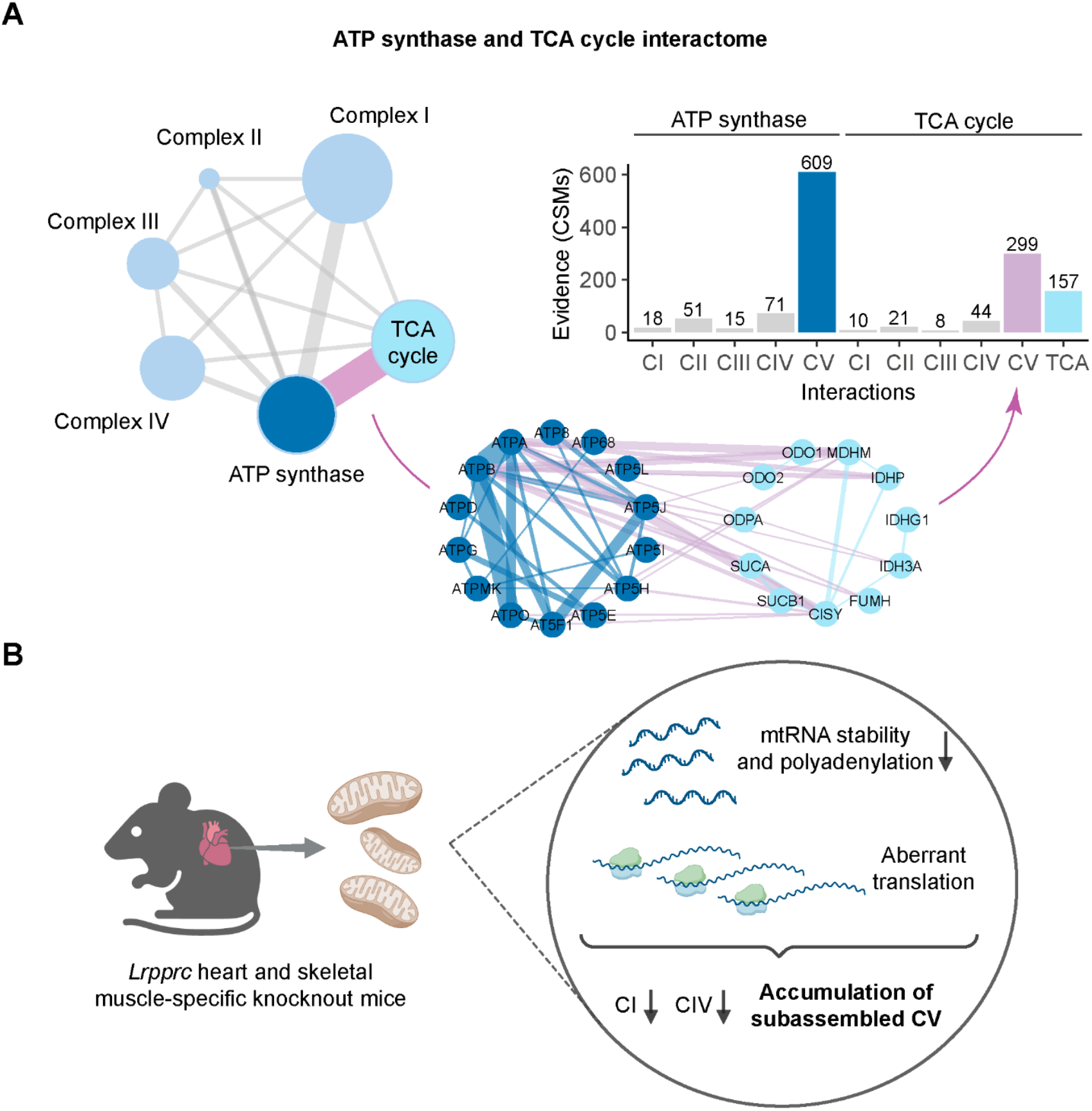
ATP synthase (complex V) interacts strongly with TCA cycle enzymes in the mouse heart. (A) Network plot illustrating the interactions between the OXPHOS complexes and the TCA cycle enzymes. Specific focus is given to the ATP synthase and TCA cycle interactions. Data represents the sum of n=3 replicates from wild-type heart mitochondria, reanalyzed data (Milenkovic et al, 2023). Thickness of the lines relate to the number of crosslinks observed. The interaction evidence was defined by the crosslink spectral matches (CSMs) shown as a histogram. (B) Schematic representation of the molecular phenotype found in the hearts of heart and skeletal muscle-specific Lrpprc knockout mice (Ruzzenente et al, 2012; Mourier et al, 2014; Kühl et al, 2017; Rubalcava-Gracia et al, 2024), abbreviations: complex I (CI), complex IV (CIV), ATP synthase (CV). Panel B was created with BioRender.com.

### Knockout of *Lrpprc* in the mouse heart perturbs the F_O_ portion of ATP synthase

Next, to investigate how the identified ATP synthase – TCA cycle interactions observed in the published dataset would be affected in mitochondrial dysfunction, we set out to perform new experiments on heart mitochondria from heart and skeletal muscle-specific *Lrppr*c knockout mice, a well-established model of impaired mitochondrial gene expression (Mourier *et al*, 2014; Ruzzenente *et al*, 2012; Kühl *et al*, 2017; Rubalcava-Gracia *et al*, 2024). LRPPRC is an RNA-binding protein required for the stabilization and translation of mitochondrial mRNAs (mtRNAs) (**Figures 1B**). In the heart, loss of LRPPRC impairs the synthesis of mtDNA-encoded OXPHOS subunits, leading to ATP synthase deficiency and accumulation of F_1_ subassemblies (Mourier *et al*, 2014; Ruzzenente *et al*, 2012; Rubalcava-Gracia *et al*, 2024). The *Lrpprc* knockout mice thus provide a robust model to investigate how ATP synthase deficiency and instability reshape its protein-interaction network.

We first performed label-free quantitative (bottom-up) proteomics on isolated heart mitochondria from wild-type and *Lrpprc* knockout mice to define global relative changes in mitochondrial protein abundance. The proteomics analyses revealed an expected profound downregulation of LRPPRC in the knockouts (**Figure 2A**). This was accompanied by a concomitant decrease in SLIRP, highlighting the essential role of LRPPRC in maintaining SLIRP stability, as previously reported (Ruzzenente *et al*, 2012; Kühl *et al*, 2017; Rubalcava-Gracia *et al*, 2024; Sasarman *et al*, 2010). Interestingly, we also observed reduced levels of several mtDNA-encoded OXPHOS subunits, including COX1, COX2 and COX3 (complex IV) and ATP6 (ATP synthase) (**Figure 2A**). Notably, the abundance of ATP8, another mtDNA-encoded subunit of ATP synthase that, together with ATP6, forms part of the membrane-embedded F_O_ domain, was found to be unchanged. In contrast, several nuclear-encoded ATP synthase subunits localized either to the F_O_ domain (ATPK, ATP5L, ATP5I) or the peripheral stalk (ATP5H, AT5F1) were upregulated, while the abundance of the catalytic F_1_ subunits remained unaffected in the LRPPRC knockout hearts (**Figure 2A**). These changes of levels of ATP synthase subunits likely represent a compensatory response to its instability, reflected by the increased expression of assembly and biogenesis factors (**Table S1**), including the ATP synthase F_1_ complex assembly factor 1 (ATPF1) (García-Bermúdez & Cuezva, 2016).

**Figure 2.**
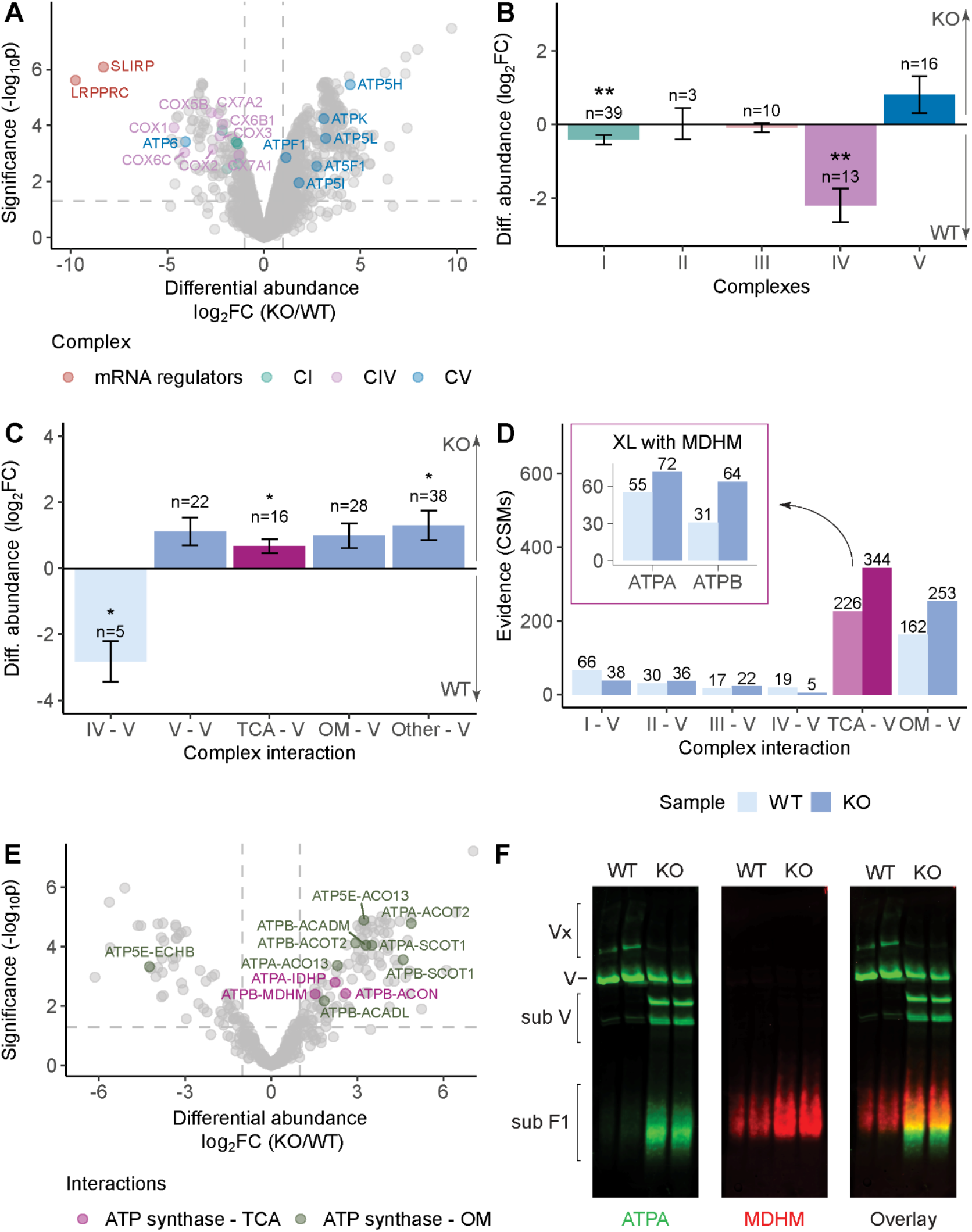
Proteomics and crosslinking analyses of wild-type and LRPPRC knockout heart mitochondria revealed increased interactions between the F1 head of ATP synthase and metabolic enzymes. (A) Differential abundance analysis of the proteins detected in the Lrpprc knockout (KO) compared to the wild-type (WT) heart mitochondria. The y-axis represents the-log10 p-value and the x-axis the log2 fold change (FC). Proteins related to mRNA regulation, complex I (CI), complex IV (CIV) and ATP synthase (CV) are highlighted with an FDR < 5% and |log2FC| >1. n=3. (B) Bar plot representing the mean differential abundance calculated as the log2 fold change of the abundance of the protein subunits of each OXPHOS complex in the proteomics data. A positive fold change indicates a higher abundance in the Lrpprc knockout compared to the wild-type heart mitochondria. n represents the number of subunits identified per complex. Number of biological replicates, n=3; error bars indicate mean ± SEM. (C) XL-MS data. Bar plot representing the mean log2 fold change of the differentially changed protein interactions between ATP synthase and other OXPHOS complexes, ATP synthase and TCA cycle proteins (TCA) and ATP synthase and other metabolic enzymes present in mitochondria (OM) or between ATP synthase and the remaining mitochondrial matrix proteins (Other), respectively. The fold change was calculated as the difference in crosslink intensity between the Lrpprc knockout and wild-type heart mitochondria. n represents the number of protein interactions identified per group. Number of biological replicates, n=3; error bars indicate mean ± SEM. (D) Evidence of protein-protein interactions calculated as the number of crosslink spectral matches (CSMs) detected for the interactions between ATP synthase and the OXPHOS complexes and metabolic proteins. The wild-type heart mitochondria are represented in light blue, and the Lrpprc knockout heart mitochondria are represented in dark blue. ATP synthase and TCA cycle interactions are highlighted in pink. A detailed view of the interactions between MDHM and ATP synthase proteins, ATPA and ATPB, is provided. Cumulative across n=3. (E) Differential abundance analysis of the interactions detected in the Lrpprc knockout compared to the wild-type heart mitochondria. The y-axis represents the-log10 p-value and the x-axis the log2 fold change. The fold change was calculated as the difference in crosslink intensity between the Lrpprc knockout and wild-type heart mitochondria. ATP synthase interactions with TCA cycle and other metabolic enzymes with an FDR < 5% and |log2FC| >1 are highlighted. (F) Fluorescent western blot of a BN-PAGE-separated wild-type and LRPPRC knockout heart mitochondria after digitonin solubilization decorated with antibodies against ATPA and MDHM and the overlay of both signals. The positions of the ATP synthase oligomers (Vx), monomers (V), subassemblies (sub V) and F1 head subassemblies (sub F1) are indicated on the left side. The partial disassembly of ATP synthase in the knockout heart mitochondria is evident, with the subassembled F1 species comigrating with MDHM, among other proteins. *Adjusted p-value < 0.01, **adjusted p-value < 0.05, ***adjusted p-value < 0.001 by Student’s t test after Benjamini-Hochberg correction (B and C).

Analyses of global OXPHOS protein levels, in which the mean fold change of all identified subunits within each complex was compared between conditions, revealed a general decrease only in complexes I and IV (**Figure 2B**) in the *Lrpprc* knockout hearts, in agreement with earlier reports (Mourier *et al*, 2014; Kühl *et al*, 2017). Beyond the OXPHOS system, proteins typically associated with mitochondrial dysfunction, including those involved in apoptosis, protein degradation, stress responses, and one-carbon metabolism, were also upregulated, consistent with previous findings by Kühl *et al*, 2017 (**Table S1**).

Moreover, we identified novel dysregulated proteins in the *Lrpprc* knockout heart mitochondria, such as components of the mitochondrial calcium uptake (MICU) complex, several solute carriers (SLCs) and tRNA aminoacylation-related enzymes (**Figure S2**).

### Increased interactions between the partially released F_1_ domain of ATP synthase and metabolic enzymes in the *Lrpprc* knockout heart mitochondria

Given the ATP synthase deficiency and structural instability in *Lrpprc* knockouts (Ruzzenente *et al*, 2012; Mourier *et al*, 2014), we next examined how these alterations affect the interaction landscape of ATP synthase. To this end, we performed in-solution XL-MS analyses on intact mitochondria isolated from wild-type and *Lrpprc* knockout hearts using an optimal concentration of 0.5 mM disuccinimidyl sulfoxide (DSSO, **Figure S3**). The crosslinking data showed high reproducibility across biological replicates (**Figure S4**). To compare the two conditions, crosslinks were quantified based on MS1 intensities and differential abundance analyses of the interactions were performed (**Table S2**). In brief, fold changes were calculated as the difference in total MS1 crosslink intensity of the interactions detected in the wild-type and LRPPRC knockout heart mitochondria. To ensure that the observed interaction changes were not confounded by protein abundance differences, we used the generated quantitative proteomics data as a reference for the evaluation of the interactomes.

We first assessed the global interaction changes among the five OXPHOS complexes, metabolic enzymes, and other proteins present in the mitochondrial matrix in the *Lrpprc* knockouts. Differential changes were detected in crosslink intensity between ATP synthase and complex IV, between ATP synthase and TCA cycle enzymes and between ATP synthase and other proteins, among which were the components of electron transfer flavoprotein system (ETFA, ETFB and ETFD), ATP synthase assembly factors and regulators of its activity (**Figure 2C, Table S2**). Interactions involving ATP synthase (subunits ATPB and ATPO) and complex IV (subunits COX6A1, COX7A2, COX5B) were reduced in the *Lrpprc* knockout mitochondria (**Figure 2C**), likely because of the overall decrease in complex IV abundance (**Figure 2A and B**). In contrast, interactions between ATP synthase and TCA cycle enzymes were markedly increased in the *Lrpprc* knockouts, as indicated by an almost twofold increase in total crosslink intensity (**Figure 2C**). This increase was further supported by the higher number of identified interactions which are represented by the crosslink spectral matches (CSMs), with 226 and 344 detected in wild-type and *Lrpprc* knockout mitochondria, respectively (**Figures 2C–D**). Loss of LRPPRC in the heart specifically enhanced crosslinking between the F_1_ head of the ATP synthase (subunits ATPA, ATPB, and ATP5E) and enzymes of the TCA cycle (ACON, IDHP, MDHM) as well as the enzymes involved in fatty-acid β-oxidation (acyl-coenzyme A thioesterase 2, ACOT2; acyl-coenzyme A thioesterase 13, ACOT13; medium-chain specific acyl-CoA dehydrogenase, ACADM, and ACADL) and ketone-body metabolism (succinyl-CoA:3-ketoacid coenzyme A transferase 1, SCOT1) (**Figure 2E**). Notably, both the F_1_ head subunits and all metabolic enzymes involved in these interactions displayed unchanged abundance in the proteomics and western blot data (**Figure S5 and S6**), showing that the elevated crosslink intensity arises from enhanced physical association between these proteins in the LRPPRC knockout hearts rather than changes in their expression levels. To validate the increased interactions in the *Lrpprc* knockouts detected by XL-MS, we examined whether subunits of the F_1_ head and TCA cycle enzymes co-migrate on Blue Native PAGE (BN-PAGE) under mild mitochondrial solubilization conditions. We focused on the F_1_ subunit ATPA and the TCA cycle enzyme MDHM, as they exhibited one of the strongest increases in crosslink intensity between the two systems (**Figure 2D**). Upon mitochondrial solubilization with digitonin, we confirmed that loss of LRPPRC led to a marked reduction in the dimeric and oligomeric ATP synthase population, consistent with impaired complex stability (**Figure 2F and S7A**). This was accompanied by the accumulation of two distinct F_1_ subassemblies containing ATPA (**Figure 2F**), as previously reported (Mourier *et al*, 2014). In addition to these subassembled forms, we detected a previously overlooked lower-molecular-weight ATPA-containing species that co-migrated with MDHM, consistent with the association detected by XL-MS (**Figure 2C-E**). Moreover, the MDHM signal on BN-PAGE appeared more intense and diffuse in *Lrpprc* knockout mitochondria, despite unchanged total MDHM protein levels (**Figures S5–S6**).

Following the BN-PAGE analyses, we also assessed ATPase *in gel* activity in the *Lrpprc* knockout mitochondria. The monomeric ATP synthase forms and the two distinct F_1_ subassemblies retained ATP hydrolytic activity, whereas the newly detected lower-molecular-weight, ATPA-containing species showed no catalytic activity (**Figure 3A**), indicating that the latter represent a partial F_1_ head. Together, these findings show that mitochondrial dysfunction caused by LRPPRC loss promotes the reorganization of MDHM into higher-molecular-weight assemblies, some of which may include partially assembled F_1_ species of ATP synthase.

**Figure 3.**
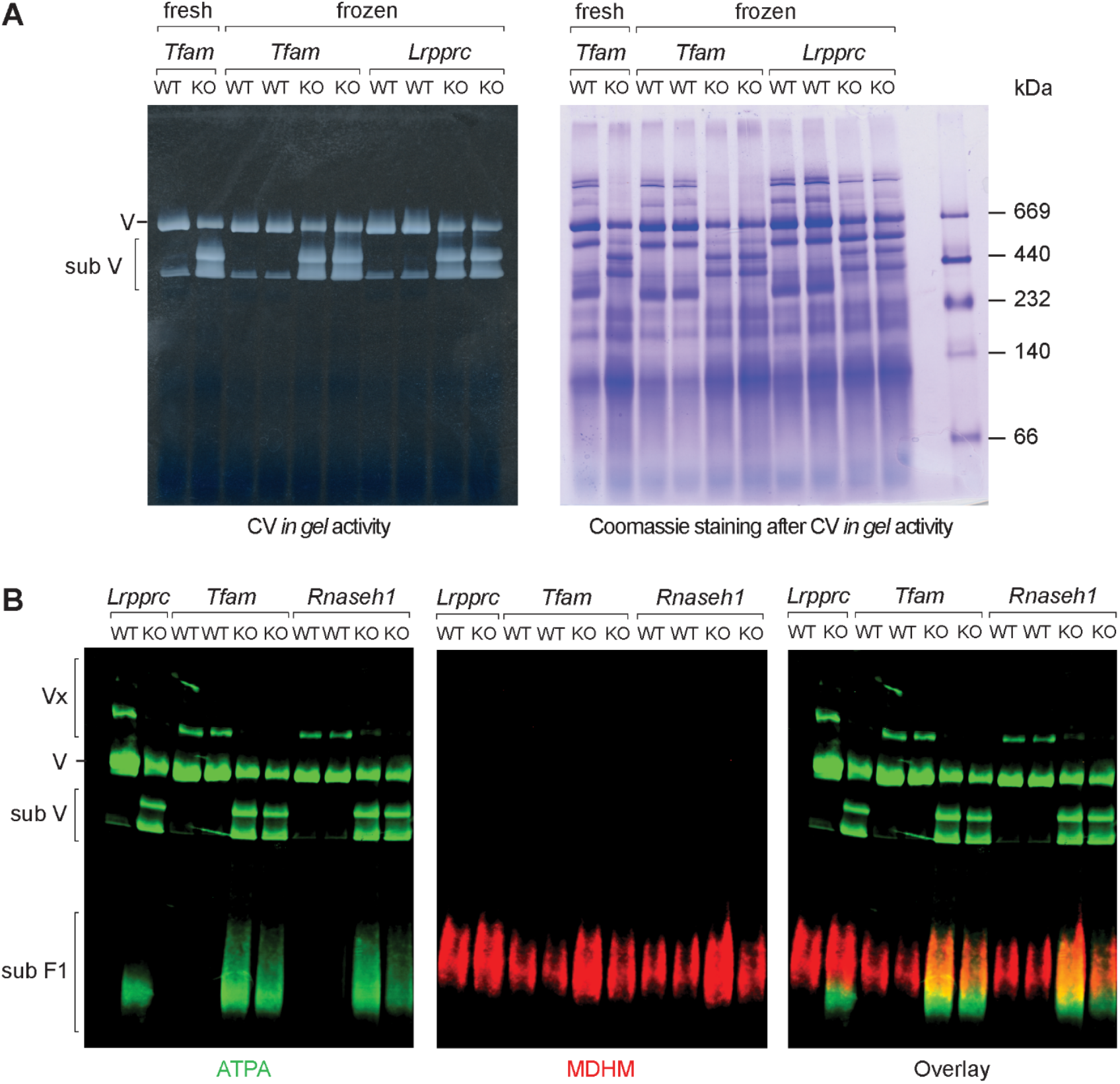
Co-migration of MDHM and ATPA in mouse models with disrupted mtDNA maintenance and expression. (A) BN-PAGE analyses of heart mitochondria followed by ATPase (CV) in gel activity show an increase in ATP synthase subassemblies in Lrpprc and Tfam knockouts (KO). No difference is observed between freshly isolated and frozen mitochondria samples after digitonin solubilization. (B) Fluorescent western blot of BN-PAGE-separated heart mitochondria from tissue-specific Lrpprc, Tfam and Rnaseh1 knockouts (KO) and their corresponding controls (WT), probed with antibodies against ATPA and MDHM and overlay of both signals. The position of the ATP synthase oligomers (Vx), monomers (V), subassemblies (sub V) and F1 head subassemblies (sub F1) is indicated on the left side (A and B).

Next, we asked whether the increased interaction between ATPA and MDHM is unique to LRPPRC deficiency or represents a more general consequence of impaired ATP synthase function and stability. To this end, we used BN-PAGE to examine heart mitochondria of other knockout mouse models affecting distinct steps of mtDNA maintenance and gene expression (**Figure 3B and S7B**). The studied mutants included heart- and skeletal muscle-specific knockouts of TFAM, a mtDNA packaging and transcription factor, and RNase H1, an enzyme required for the initiation and completion of mtDNA replication (Larsson *et al*, 1998; Misic *et al*, 2022). Similar to the *Lrpprc* knockout, additional lower-molecular-weight ATPA-containing species co-migrated with MDHM, which displayed a stronger and more diffuse signal in both *Tfam* and *Rnaseh1* knockout mitochondria (**Figure 3A**). These findings show that the enhanced association between ATP synthase and metabolic enzymes likely represents a common response to ATP synthase dysfunction and instability.

### LRPPRC loss induces ATIF1-mediated regulatory remodeling at the catalytic F_1_ domain of ATP synthase

We next examined interactions between ATP synthase subunits and suggested modulators of its activity. Among the dysregulated associations, we observed increased interlinks with PPIF (Cyclophilin D) and ATIF1 in the *Lrpprc* knockout mitochondria. Interactions between the ATPB subunit of the F_1_ head and PPIF were increased (**Figure S8**). The precise mechanism by which PPIF exerts its regulatory effect on ATP synthase remains unknown (Porter & Beutner, 2018). Although an interaction between PPIF and OSCP (ATPO) has been reported biochemically (Giorgio *et al*, 2009), we did not observe OSCP crosslinks in our XL-MS dataset. However, we detected PPIF crosslinks with the ATPA and ATPB subunits of ATP synthase. Importantly, PPIF-ATPB interactions were increased in LRPPRC knockout mitochondria, consistent with stress-induced remodeling of the F_1_ head and an enhanced susceptibility to permeability transition pore (PTP) activation. ATIF1, in contrast, acts as a regulatory protein with a dual role in stabilizing ATP synthase dimers and modulating the catalytic activity of the F_1_ domain (Esparza-Moltó *et al*, 2017). In the absence of LRPPRC, we observed a vast increase in the detected crosslinks of ATIF1 to the F_1_ head of ATP synthase (**Figure 4A and 4B**).

**Figure 4.**
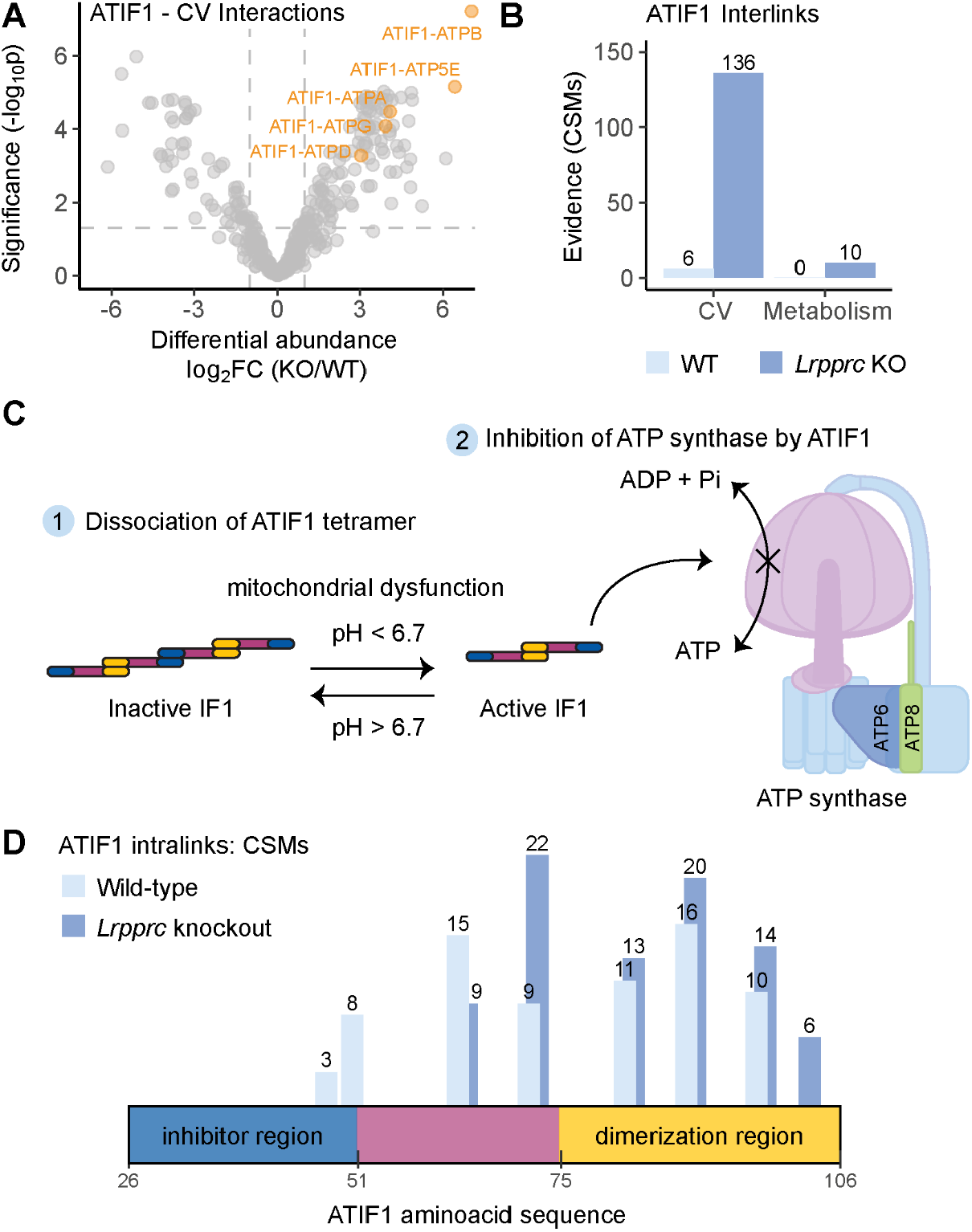
ATIF1 dissociates into dimers upon mitochondrial dysfunction to inhibit the ATP synthase. (A) Differential abundance analysis of the interactions between ATP synthase and ATIF1 as detected by XL-MS analyses of Lrpprc knockout (KO) and wild-type (WT) heart mitochondria. The y-axis represents the-log10 p-value and the x-axis the log2 fold change (FC). The fold change was calculated as the difference in crosslink intensity between the Lrpprc knockout and wild-type heart mitochondria. ATP synthase (CV) interactions with metabolic proteins with an FDR < 5% and |log2FC| >1 are highlighted. (B) Evidence of protein interactions represented by the cross¬link spectral matches (CSMs) detected in the WT (light blue) and the Lrpprc knockout (dark blue) samples showing interactions between ATIF1 and ATP synthase or proteins related to mitochondrial metabolism in the matrix. Cumulative across n=3 biological replicates. (C) Schematic representation of the dissociation of the ATIF1 tetramer into dimers, followed by binding and inhibition of the ATP synthase. The F1 region of the ATP synthase is colored in pink. (D) Distribution of ATIF1 intralinks across the protein sequence, calculated as CSMs for the wild-type (light blue) and Lrpprc knockout (dark blue) samples. (D) Crosslink spectral matches (CSMs) detected for the interaction between ATIF1 proteins in the wild-type (light blue) and the Lrpprc knockout (dark blue) samples per region. The ATIF1 regions were defined based on their functional properties: inhibition (residue 26-52), dimerization (residue 74-106), and other (residue 53-63). Cumulative across n=3 biological replicates.

Previous *in vivo* studies in cells and mice have shown that ATIF1 itself exists in inactive tetrameric or oligomeric forms, as well as in an active dimeric form that binds to ATP synthase (Romero-Carramiñana *et al*, 2023; García-Bermúdez & Cuezva, 2016; Weissert *et al*, 2021; Esparza-Moltó *et al*, 2017). In *Lrpprc* knockout mitochondria, ATP synthase dysfunction and the accumulation of F_1_ subassembled species account for the substantially increased crosslink intensity observed between ATIF1 and the F_1_ head. These findings are consistent with cryo-EM data showing that ATIF1 exerts its inhibitory effect through direct binding to the F_1_ catalytic domain (Romero-Carramiñana *et al*, 2023; Gu *et al*, 2019; Cabezón *et al*, 2003). To further investigate the structural basis of ATIF1 binding to ATP synthase in mitochondria under native conditions, we examined the conformational differences between its active and inactive forms. ATIF1 can undergo pH-dependent oligomeric rearrangements (Boreikaite *et al*, 2019). At physiological pH values above 6.7, it predominantly exists in an inactive tetrameric or oligomeric state, composed of two or more dimers linked via the N-terminal inhibitory region (**Figure 4C**). When respiratory chain function is compromised, as in the *Lrpprc* knockout hearts, the proton gradient collapses and the pH within the mitochondrial matrix can drop below 6.7, triggering the dissociation of ATIF1 oligomers into dimers. This transition exposes the N-terminal inhibitory region, allowing ATIF1 to bind the F_1_ domain and inhibit its catalytic activity (**Figure 4C**) (Gu *et al*, 2019). Consistent with this model, intralinks detected within ATIF1 in wild-type mitochondria spanned the entire protein sequence (**Figure 4D**), indicating that both the N-terminal inhibitory and C-terminal dimerization regions participate in intermolecular interactions within ATIF1 oligomers. In contrast, in *Lrpprc* knockout mitochondria, the intralink distribution shifted substantially toward the C-terminal dimerization region, leaving the N-terminal inhibitory region available for binding to the F_1_ domain of ATP synthase to suppress its catalytic activity (**Figure 4D**). This interpretation is further supported by the distribution of the interlinks across the ATIF1 protein sequence. In the absence of LRPPRC, ATIF1 no longer forms higher-order oligomers but instead uses its N-terminal inhibitor region to bind directly to the ATP synthase subunits ATPA, ATPB, and ATPG (**Figures 4B and S9**). The interaction between the N-terminal inhibitory region of ATIF1 and ATP synthase was not detected in wild-type mitochondria, suggesting that this binding occurs uniquely under conditions of ATP synthase dysfunction or instability to regulate its reverse, ATP-hydrolytic mode.

Additionally, interlinks between the C-terminal dimerization region of ATIF1 and metabolic enzymes such as MDHM, SDHA, and CSIY were also detected, providing additional evidence for their increased association with ATP synthase in *Lrpprc* knockout mitochondria (**Figures 2C–E**). Together, these findings demonstrate that in *Lrpprc* knockout hearts, both ATIF1-mediated inhibition and the broader interactome remodeling occur within the F_1_ catalytic domain of ATP synthase.

### Structural reorganization of ATP synthase in the Lrpprc knockout hearts

Beyond mapping interaction changes, the in situ XL-MS dataset also allowed us to examine detailed structural rearrangements within ATP synthase in the absence of LRPPRC. Analyses of intralinks within the ATP synthase revealed no significant changes among the F_1_ subunits. In contrast, loss of LRPPRC led to an increase in intralinks of the ATP5I and ATP5L subunits (**Figure 5**), which are known to play a key role in ATP synthase dimerization (Gu *et al*, 2019). The elevated intralinks of the F_O_ membrane domain likely reflect strengthened interactions between the ATP5I and ATP5L subunits of adjacent ATP synthase monomers, possibly acting as a compensatory mechanism to counteract dimer instability (**Figure 2F**). Alternatively, the increased intralinks within these subunits could result from structural rearrangements within the F_O_ region caused by ATP6 downregulation (**Figure 1A**). However, as both ATP5I and ATP5L were upregulated in the bottom-up proteomics data (**Figure 2A**), the observed increase in intralinks could also be a consequence of higher protein abundance rather than being caused by structural remodeling. Furthermore, analyses of the interlinks involving ATP synthase subunits whose abundance was unchanged in bottom-up proteomics revealed increased crosslinks between F_O_ membrane subunit ATP8 and the peripheral stalk subunit ATP5J, accompanied by decreased crosslinks between F_1_ head subunit ATPB and the same peripheral stalk subunit in the absence of LRPPRC (**Figure 5**). These changes indicate strengthened interactions between the F_O_ and peripheral stalk regions and weakened contacts between the F_1_ head and the peripheral stalk, in line with partial detachment of the F_1_ domain from the remaining ATP synthase structure in the *Lrpprc* knockout hearts (**Figure 2F, 3A-B**) (Mourier *et al*, 2014). Collectively, our data show that loss of LRPPRC leads to reduced ATP6 levels, which in turn perturbs the F_O_ region of ATP synthase. This perturbation appears to trigger a compensatory increase in other F_O_ subunits, strengthening their interactions with each other and the peripheral stalk, but weakening the coupling between the F_1_ head and the rest of the complex. Together, these changes destabilize the ATP synthase and lead to the emergence of subassembled F_1_ species.

**Figure 5.**
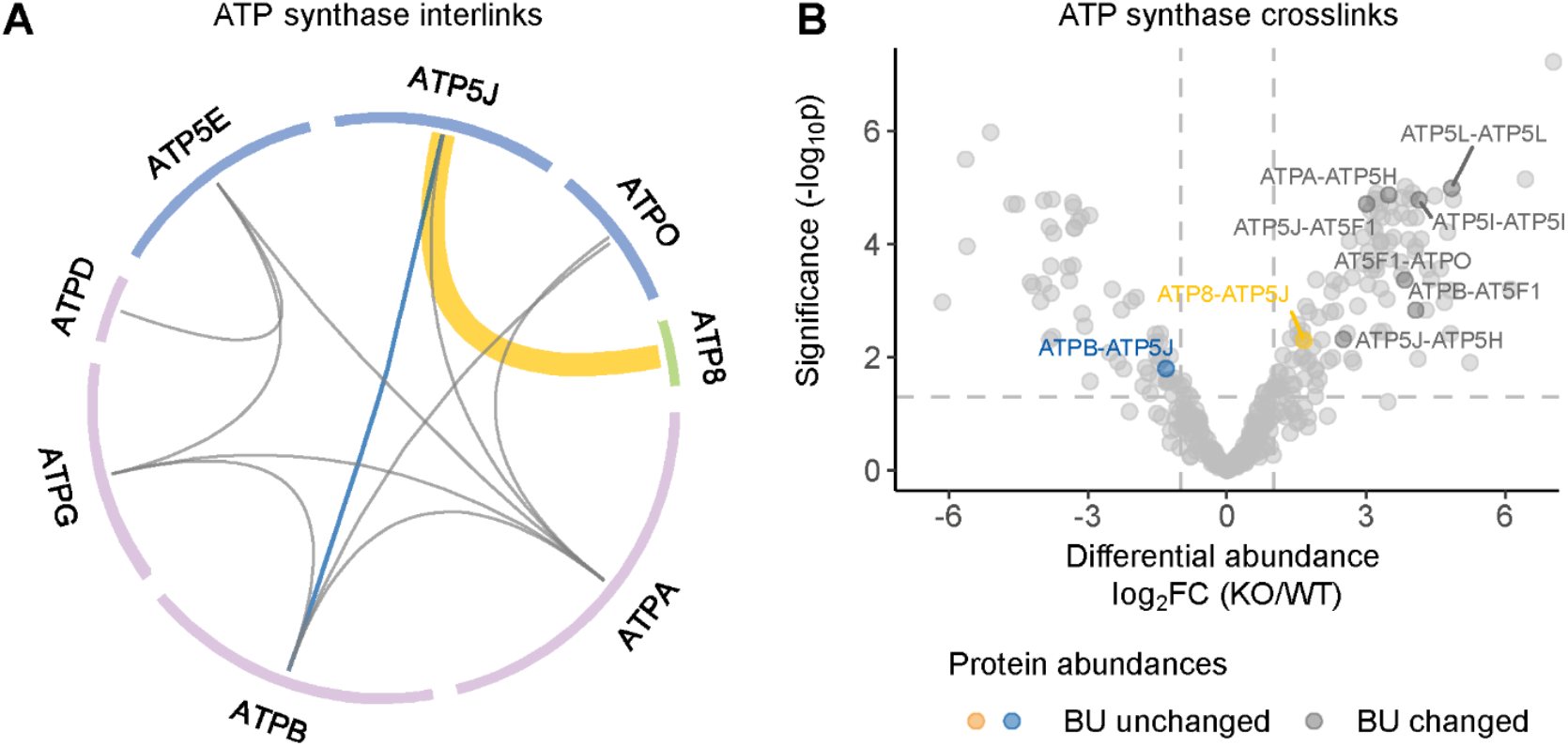
Changes in ATP synthase inter- and intralinks upon loss of Lrpprc in the heart. (A) Chord diagram representing the observed interlinks within the ATP synthase. The interactions significantly upregulated in the Lrpprc knockout are colored in yellow, and those downregulated are colored in blue. The thickness of the line is proportional to the observed fold change. Proteins are color-coded based on their location in the ATP synthase structure: F1 domain (pink), FO domain (blue) and peripheral stalk (green). (B) Differential abundance analysis of the ATP synthase crosslinks detected by XL-MS in mitochondria of Lrpprc knockout hearts (KO) in comparison with wild-type (WT) hearts. The y-axis represents the-log10 p-value and the x-axis the log2 fold change (FC). The fold change was calculated as the difference in crosslink intensity between the Lrpprc knockout and wild-type heart mitochondria. Crosslinks involving proteins without significantly changed protein abundances according to the bottom-up (BU) proteomics analysis are highlighted in yellow and blue. Crosslinks colored in grey are between subunits that also have increased protein abundances according to the BU proteomics analysis. The annotated crosslinks have an FDR < 5% and |log2FC| >1. Number of biological replicates, n=3.

## Discussion

Mitochondrial ATP synthase is well recognized for its dual role in driving ATP production and maintaining cristae architecture. Here, we show that it is not merely a structural and catalytic component of the OXPHOS system but also acts as a dynamic hub that links mitochondrial bioenergetics with central carbon metabolism. Using *in situ* XL-MS on intact mitochondria combined with quantitative proteomics and BN-PAGE analyses, we demonstrate that ATP synthase directly associates with enzymes of the TCA cycle in the mouse heart and undergoes extensive structural, regulatory, and interactome remodeling when its function and stability are compromised.

Under normal physiological conditions, the F_1_ catalytic head of ATP synthase forms extensive contacts with multiple TCA enzymes, including citrate synthase, isocitrate dehydrogenases, oxoglutarate dehydrogenase, succinyl-CoA ligase, fumarate hydratase, and malate dehydrogenase within intact heart mitochondria, (**Figure 1A, S1**). This finding establishes a previously unappreciated link between the OXPHOS and the TCA cycle that goes beyond complex II, which has been considered to be the sole bridge between the two pathways. Of note, cross-species reanalyses of XL-MS data, including boar sperm, bovine heart, and human HEK293T mitochondria, confirmed that the F_1_ catalytic head of ATP synthase universally forms extensive contacts with multiple TCA enzymes (Leung *et al*, 2021; Hevler *et al*, 2021; Zhu *et al*, 2024). All these identified interactions may enhance both the efficiency and regulation of mitochondrial energy transduction, to allow high-energy-demand tissues such as the heart to dynamically coordinate ATP synthesis with metabolic flux in real time. Consistent with this idea, *in vitro* studies have shown that TCA cycle enzymes can assemble into dynamic metabolon-like structures that mediate substrate channeling and reorganize in response to substrate availability and the metabolic state (Bulutoglu *et al*, 2016). The physiological significance of these assemblies remains debated, with a recent review proposing that they may form transiently to optimize metabolic efficiency under conditions of high energy demand (Omini *et al*, 2024). In support of such functional coupling, MDHM-generated oxaloacetate is known to inhibit complex II of the respiratory chain to regulate flux between complexes I and II (Molinié *et al*, 2022). Our findings suggest that the ATP synthase itself could act as a structural and functional component of such larger metabolon-like networks in cardiac mitochondria, thereby linking local metabolic flux directly to ATP synthesis. When mitochondrial gene expression is disrupted, as in the heart- and skeletal-muscle-specific *Lrpprc* knockout model used here, the ATP synthase undergoes a profound structural remodeling (**Figure 5**) that ultimately perturbs the cristae morphology (Mourier *et al*, 2014) (**Figure 6**). The selective downregulation of mtDNA-encoded subunits, notably ATP6, destabilizes the F_O_ domain of ATP synthase, triggering compensatory upregulation of other F_O_ subunits (ATP5I and ATP5L) and strengthening the interactions between them (**Figure 2A and 5**). In contrast, F_1_ and peripheral stalk subunits remain unchanged or are even moderately upregulated (**Figure 2A**). We also detected stronger interactions between F_O_ and the peripheral stalk, accompanied by weakened association between the F_1_ head and the rest of the complex. The observed compositional imbalance, together with altered intra- and inter-subunit contacts, promotes partial disassembly of the complex and accumulation of soluble F_1_ subcomplexes in the *Lrpprc* knockouts (**Figure 6**). These findings are consistent with a defect in the late stages of ATP synthase assembly, during which the incorporation of mtDNA-encoded ATP6 and ATP8 into the F_O_ membrane domain is required to anchor the F_1_ head and stabilize the dimeric form of the complex (Spikes *et al*, 2020). Similarly, patients carrying mutations in ATP6 or ATP8 often display loss of dimeric ATP synthase forms and accumulation of F1 subspecies (Jonckheere *et al*, 2008; Jackson *et al*, 2017; Dautant *et al*, 2018). Previous studies have shown that the incorporation of peripheral stalk subunit ATP5J precedes the integration of ATP6 and ATP8 (He *et al*, 2020). In the *Lrpprc* knockout hearts, ATP5J upregulation and its strengthened interaction with ATP8 (**Figure 5**) likely represent an adaptive adjustment of the assembly pathway to partially stabilize the F_O_ domain in response to ATP6 deficiency. Together, the reinforced interactions observed among some F_O_ subunits and their increased contacts with the peripheral stalk may serve to preserve partial structural integrity and prevent complete dimer dissociation in the absence of LRPPRC. Meanwhile, the released F_1_ subcomplexes engage in enhanced interactions with the enzymes of the TCA cycle, fatty-acid β-oxidation, and ketone-body metabolism, as revealed by XL-MS (**Figure 2E, Table S2**).

**Figure 6.**
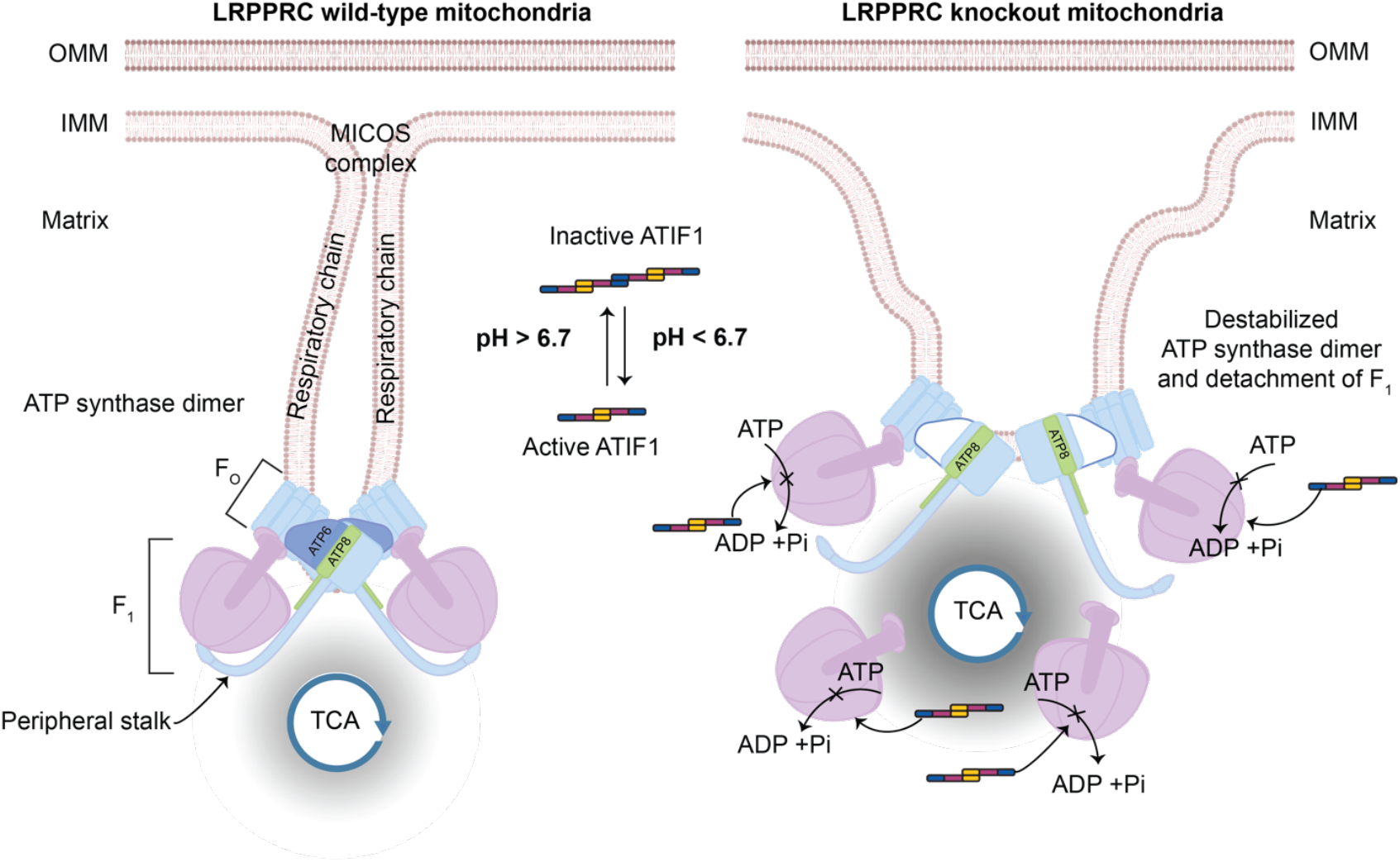
Graphical model of ATP synthase organization, its interactome and its regulatory remodeling in wild-type and LRPPRC knockout heart mitochondria. In wild-type heart mitochondria, ATP synthase dimers are positioned at the tips of inner mitochondrial membrane (IMM) cristae. The complex consists of the membrane-embedded F_0_ domain, which contains the only mtDNA-encoded subunits (ATP6 and ATP8) and the matrix-exposed F_1_ catalytic head, connected via the peripheral stalk. In LRPPRC knockout heart mitochondria, reduced ATP6 levels trigger compensatory rearrangements within the F_0_ domain that ultimately destabilize the entire complex and promote partial detachment of the F_1_ head. This structural destabilization, in turn, perturbs cristae membrane organization (Mourier et al, 2014). Under physiological conditions, the endogenous ATP synthase inhibitor (ATIF1) exists predominantly as tetramers or higher-order oligomers formed by N-terminally mediated dimer-dimer interactions. In LRPPRC-deficient heart mitochondria, matrix acidification caused by respiratory chain dysfunction promotes the dissociation of ATIF1 oligomers into active dimers, exposing the N terminal-inhibitory domain, which then binds to the F_1_ catalytic domain and inhibits ATP hydrolysis. The TCA cycle is depicted in proximity to ATP synthase, with light gray shading indicating the prominent crosslinks identified between the F_1_ head and TCA cycle enzymes in wild-type mitochondria. Darker gray shading represents the further increase in crosslink abundance among the F_1_ head, ATIF1, and TCA cycle enzymes in LRPPRC knockout heart mitochondria as compared to the wild type. For completeness, the location of the MICOS complex is shown in wild-type mitochondria, as well as the location of respiratory chain complexes along the planar regions of cristae membranes. Abbreviations: OMM-outer mitochondrial membrane.

Our BN-PAGE analyses further corroborate the *in situ* XL-MS findings, confirming the reorganization of ATP synthase and its altered interactions in the *Lrpprc* knockout hearts (**Figures 2F and 3**). The F_1_ subunit ATPA co-migrate with MDHM in *Lrpprc* knockouts as well as in other models of impaired mtDNA maintenance and expression in the heart, such as *Tfam* and *Rnaseh1* knockouts. These findings suggest that the enhanced association between the F_1_ head and metabolic enzymes, exemplified here by MDHM, represents a general feature that emerges under conditions of defective mtDNA expression and subsequent ATP synthase dysfunction. Further supporting our data, a complexome profiling study of wild-type heart mitochondria also detected co-migration of ATPA, ATPB, MDHM, and CISY at comparable molecular weights (reanalyzed data in **Figure S10**) (Yin *et al*, 2024), indicating that these associations are present at a basal level under normal physiological conditions and are therefore captured in our XL-MS dataset of wild-type hearts (**Figure 1 and S1**). In our BN-PAGE analyses, co-migration became detectable only under conditions of ATP synthase dysfunction, likely because the interactions become more abundant upon structural destabilization of the enzyme, as evidenced by the increased crosslink intensities observed in the *Lrpprc* knockout hearts (**Figure 2C-E**).

We also identify a regulatory remodeling of ATP synthase mediated by ATIF1. The *Lrpprc* knockout mitochondria, yielded increased crosslinks of ATIF1 to the F_1_ head, consistent with its activation under sub-physiological pH conditions, when the proton gradient collapses (Cabezon *et al*, 2000; Boreikaite *et al*, 2019) (**Figure 6**). Our XL-MS data provide *in vivo* evidence that ATIF1 transitions from inactive oligomers to inhibitory dimers upon activation, which leads to the N-terminal inhibitory region engaging with the catalytic subunits of F_1_ domain in *Lrpprc* knockouts. This conformational switch prevents wasteful ATP hydrolysis by effectively shifting the enzyme from an energy-producing to an energy-preserving state. Our findings are consistent with previous reports showing that ATIF1 activity is also modulated by PKA-dependent phosphorylation at Ser39 within the N-inhibitory domain. Phosphorylation at this site prevents ATIF1-binding to F_1_ under normal conditions (García-Bermúdez *et al*, 2015). Notably, we also detect crosslinks between ATIF1 and TCA cycle enzymes in the *Lrpprc* knockouts (**Figure S9**) suggesting that ATP synthase inhibition is integrated with a broader metabolic remodeling program in mitochondria.

In summary, our findings reveal a coordinated structural and interactome remodeling of ATP synthase that integrates metabolic, architectural, and inhibitory mechanisms. Under normal conditions, ATP synthase resides at the interface between the OXPHOS and the TCA cycle, ensuring efficient coupling of energy conversion with metabolic flux. Under mitochondrial stress, the partly disassembled enzyme associates with metabolic enzymes and recruits its intrinsic inhibitor to adopt an energy-preserving state. This remodeling may hold broader relevance for mitochondrial pathophysiology. Defects in ATP synthase assembly or mtDNA expression underlie a spectrum of human disorders, including mitochondrial encephalomyopathies, cardiomyopathies, and neurodegenerative diseases (Galber *et al*, 2021). The enhanced association of ATP synthase and metabolic enzymes observed under mitochondrial dysfunction may represent an adaptive mechanism that sustains local energy homeostasis when oxidative phosphorylation fails. Future studies dissecting this remodeling process may provide new insight into how cells withstand bioenergetic failure and identify potential therapeutic strategies for conditions involving impaired ATP synthase activity or stability.

## Methods

### Animals and housing

Heart- and skeletal muscle–specific *Lrpprc* knockout mice on a C57BL/6N background were generated as described earlier (Ruzzenente *et al*, 2012). *Tfam* and *Rnaseh1* knockout mice with the same tissue specificity and background were generated as reported earlier (Larsson *et al*, 1998; Misic *et al*, 2022). For experiments, mice were euthanized by cervical dislocation, and hearts were collected at the latest viable time points (10–12 weeks of age for *Lrpprc* knockouts, 7–9 weeks of age for *Tfam* knockouts and 24 weeks for *Rnaseh1* knockouts). Animals were maintained in individually ventilated cages under a 12-h light/dark cycle with ad libitum access to chow and water. All procedures were approved by the local animal ethics committee and carried out in accordance with national and European legislation.

### Mitochondria isolation

For *in situ* crosslinking, intact mitochondria were prepared in isolation buffer (MIB) containing 200 mM sucrose, 20 mM HEPES, and 1 mM EDTA. For all other purposes, mitochondria were prepared in MIB consisting of 235 mM sucrose, 20 mM Tris-HCl (pH 7.4), and 1 mM EDTA. Hearts were excised after cervical dislocation and transferred into ice-cold PBS. After rinsing to remove blood, hearts were minced on ice and transferred into 1.8 Ml of MIB supplemented with 200 µL of 2.5% trypsin. Samples were rotated at 4 °C for 10 min, diluted with MIB containing 0.04% trypsin inhibitor, and homogenized. The homogenate was centrifuged at 1,000 × g for 10 min at 4 °C, and the supernatant was collected and centrifuged at 10,000 × g for 10 min. The resulting pellet was resuspended in MIB containing 0.2% BSA, adjusted to 12 mL, gently mixed, and centrifuged again at 10,000 × g. The final pellet was washed twice with MIB lacking BSA and trypsin inhibitor, then resuspended in 150 µL of MIB. All steps were carried out on ice or at 4 °C. Mitochondrial protein concentration was determined using a Qubit fluorometer (Invitrogen).

### BN-PAGE

BN-PAGE was performed as previously described (Milenkovic *et al*, 2023). Mitochondrial proteins (75 µg) were incubated on ice in 50 µL of buffer containing 20 mM Tris-HCl (pH 7.4), 50 mM NaCl, 0.1 mM EDTA, and 10% (v/v) glycerol. Proteins were solubilized with 1–3% (w/v) digitonin (Calbiochem), after which native loading dye (5% Coomassie Brilliant Blue G-250, 150 mM Bis-Tris, 500 mM ε-aminocaproic acid, pH 7.0) was added. Samples were subsequently loaded onto in-house prepared 3–13% gradient BN-PAGE gels for separation of protein complexes.

### ATPase *in gel* activity assay

To visualize ATPase activity, BN-PAGE gels were incubated in a solution containing 50 mM glycine, 5 mM MgCl_2_, 0.1% Triton X-100, and 0.5 mg/mL lead nitrate, adjusted to pH 8.4. After an initial 30 min incubation in 50 mL of this buffer, an additional 50 mL of the same buffer supplemented with 4 mM ATP was added, followed by a 1 h incubation. The gels were then rinsed with water and scanned against a dark background.

### Western blotting

Proteins separated by SDS-PAGE or BN-PAGE were transferred to Hybond polyvinylidene fluoride (PVDF; GE Healthcare) membranes. For fluorescence-based detection, membranes were blocked in Intercept Blocking Buffer (LI-COR), incubated with primary antibodies, and probed with donkey anti-mouse IgG 800CW and goat anti-rabbit IgG 680RD secondary antibodies (LI-COR). Fluorescent signals were visualized using the LI-COR Odyssey imaging system. Chemiluminescent detection was performed using Clarity Western ECL Substrate (Bio-Rad). Primary antibodies included anti-ATP5A (Abcam, ab14748), anti-MDH2 (Abcam, ab96193), anti-SDHA (Abcam, ab14715), and anti-CS (Santa Cruz, sc-390693).

### Optimization of crosslinker concentration

The concentration of DSSO was optimized as previously described (Klykov *et al*, 2018). Briefly, intact mouse heart mitochondria purified from three biological replicates of wild-type and LRPPRC knockout mice were treated with 0.25, 0.5, or 1 mM DSSO for 45 min at 15°C and compared to control samples incubated at either 4 or 15°C. Subsequently, the reaction was quenched for 30 min at 15°C, and the crosslinked samples were visualized by SDS-PAGE.

### Crosslinking of mouse heart mitochondria

Intact mouse heart mitochondria purified from three biological replicates of wild-type and LRPPRC knockout mice were treated with 0.5 mM DSSO for 45 min at 15°C, followed by quenching for 30 min at 15°C. Samples were processed as previously described by Milenkovic *et al*, 2023. Briefly, crosslinked mitochondria were pelleted at 11,000 x g at 4 °C for 10 min and resuspended in 5 times the volume of lysis buffer (7 M Urea, 1% [v/v] Triton-X-100, 100 mM Tris pH 8.5, 5 mM tris(2-carboxyethyl)phosphine (TCEP), 30 mM chloroacetamide (CAA), proteinase inhibitor cocktail, 1% [vol/vol] benzonase and 2 mM Mg^2+^). Mitochondria were solubilized for 30 min on ice and centrifuged at 18,000 x g for 30 min. The supernatant was collected, 1% benzonase (v/v) was added and the sample was incubated for 2 h at room temperature while shaking. The mitochondrial proteins were precipitated following the Methanol/Chloroform precipitation protocol as previously described (Wessel & Flügge, 1984). The dry protein pellet was resuspended in digestion buffer (1% sodium deoxycholate [w/v], 100 mM Tris pH 8.5, 5 mM TCEP and 30 mM CAA) to a final protein concentration of 1µg/µL. Protein digestion was performed first with LysC (1:100 enzyme to protein ratio) for 45 min at 37°C, followed by overnight digestion with trypsin (1:25 enzyme to protein ratio) at 37°C. Digestion was stopped by addition of trifluoroacetic acid (TFA) to a final concentration of 0.5% (v/v) TFA. Samples were centrifuged at 18,000 x g at 4 °C for 10 min and the supernatant was collected.

Finally, peptides were desalted using solid-phase extraction C18 columns (Sep-Pak, Waters) and fractionated into 27 fractions for DSSO using an Agilent 1200 HPLC pump system (Agilent) coupled to a strong cation exchange (SCX) separation column (Luna SCX 5 mm to 100 Å particles, 50 x 2 mm, Phenomenex). The fractions were desalted using Oasis.

### LC-MS/MS analysis

The fractionated samples were analyzed using a Exploris 480 (ThermoFisher) mass spectrometer coupled to an Ultimate3000 UHPLC system (ThermoFisher). The fractions were injected at 3 µL/min for 1 min into a trap column (Acclaim Pepmap 100 C18, 5 mm x 0.3 mm, 5 µm, Thermo Fisher Scientific). The peptide separation was performed in a 50 cm long analytical column with a 75 µm inner diameter packed in house with C18 beads (Reprosil C18, 1.9 µm) at a column temperature of 32°C. Peptides were eluted at 300 nL/min with a total run time of 70 min, using a gradient with solvent A (0.1 % (v/v) formic acid) and solvent B (80 % (v/v) acetonitrile and 0.1 % (v/v) formic acid). Gradient separation on the analytical column: 9 % B for 1 min, from 9 % to 13 % B in 1 min, from 13 % to 41 % in 55 min, from 41 % to 99 % in 1 min, 99 %B for 4 min, 99 % to 9 % in 1 min and 9 % B for 7 min. The MS acquisition method was a data-dependent acquisition mode using the Orbitrap analyzer at 60K mass resolution in the scan range 375-2200 m/z, with standard AGC target and automatic maximum injection time. Ions with charges between +3 and +8 were selected for MS2 and filtered based on dynamic exclusion with a mass tolerance of 10 ppm and mass resolution of 30K. Peptides were fragmented with stepped HCD of 19, 25 and 28 %.

### MS Data analysis

For proteomics analysis the raw files corresponding to all fractions were analyzed with MaxQuant (version 2.4.14.0) using the full *Mus musculus* reference proteome (downloaded from UniProt 15/02/2024 with UniProtID: UP000000589). A FASTA file containing all identified proteins across both XL-MS experiments was generated, the mitochondrial transit peptides were annotated with TargetP 2.0 (Armenteros *et al*, 2019) and subsequently removed. For the DSSO XL-MS experiments, data were analyzed in Proteome Discoverer with the XlinkX node for analysis of crosslinked peptides, as previously reported (Klykov *et al*, 2018). For the XlinkX search tryptic digestion with a maximum of three misscleavages, 10ppm error for MS1 and 20ppm error for MS2. Carbamidomethyl (C) was set as a static modification, and oxidation (M) and protein N-terminal acetylation as variable modifications. Crosslinked peptides were accepted with XlinkX score higher or equal to 40 and a maximum FDR of 5%. Cros-links present in less than 2 replicates per sample were excluded. Crosslinked peptides were quantified by the addition of the MS1 precursor intensities across all spectral matches. Crosslinks without precursor intensities were excluded for analysis.

### Statistical analysis of proteomics and crosslinking data

Data analysis and visualization were performed with custom R scripts. Data were analyzed in R version 4.4.0 running in RStudio 2024.12.1 (Build 563). Data handling and visualization were mostly performed with *tidyverse* (v2.0.0), *protti* (v0.8.0), and *cowplot* (v1.1.3). Briefly, precursor intensities of the proteins or of the crosslinks were median normalized and log^2^ transformed. Imputation was performed based on the “Ludovic” method, followed by differential abundance analysis with a “moderated t-test” using the *protti* package in R. Differential abundance of the proteins and protein-protein crosslinks was defined by adjusted pvalue < 0.05 and |log^2^ fold change| >1.

## Data availability

The proteomics and crosslinking mass spectrometry data have been deposited in the ProteomeXchange Consortium via PRIDE (Perez-Riverol *et al*, 2025) partner repository with the dataset identifier PXD071441.

## Author contributions

Conceptualization: L.P.P., J.M., N.-G.L., A.J.R.H.; Data curation: L.P.P., J.M.; Formal analysis: L.P.P., J.M.; Funding acquisition: N.-G.L., A.J.R.H.; Investigation: L.P.P., J.M.; Methodology: L.P.P., J.M., T.K.; Supervision: N.-G.L., A.J.R.H.; Visualization: L.P.P., J.M.; Writing - original draft: L.P.P., J.M., N.-G.L., A.J.R.H.; and Writing- review & editing: all authors.

## Acknowledgements

We would like to thank Alisa Potter, Utrecht University, for her help in complexome profiling data analysis. N.-G.L. was supported by the Swedish Research Council (2015-00418), the Swedish Cancer Foundation (243520Pj), the Knut and Alice Wallenberg Foundation (2023.0224 and 2024.0081), the Swedish Brain Foundation (FO2025-0061-HK-269), the European Research Council (Advanced Grant 101141290), the Novo Nordisk Foundation (NNF25OC0105033), and by grants provided by Region of Stockholm (ALF project). A.J.R.H. was supported by the Netherlands Organization for Scientific Research (NWO) through the Spinoza Award (SPI.2017.028), and the European Research Council (Advanced Grant REVAMP 101141457).

## Supplementary Information

The supplementary information file contains additional Figures S1-S10, as referenced in the text. Supplementary Tables S1 and S2, as referenced in the text, are provided and include the differential abundance analysis results for the bottom-up proteomics and XL-MS experiments, respectively.

