## Supplemental Information for "Uncovering the link between ATP synthase and the TCA cycle by crosslinking mass spectrometry"

### Contents

#### Figures

1. XL-MS provides evidence for extensive interactions within and between the OXPHOS and TCA cycle proteins in mouse heart mitochondria.
2. Volcano plot highlighting previously undetected dysregulated mitochondrial proteins in heart mitochondria of *Lrpprc* knockout mice.
3. SDS-PAGE used for the optimization of crosslinker concentration for the XL-MS experiments on mouse heart mitochondria.
4. Overview of CSMs detected per replicate in the XL-MS experiments.
5. $Log_2$ fold change (FC) of mitochondrial metabolism proteins up-regulated in the interactions with ATP synthase in the Lrpprc knockout hearts.
6. Western blot analyses of SDS-PAGE-separated heart mitochondria from Lrpprc and Tfam knockout (KO) and their corresponding controls (WT).
7. Loading control for the fluorescent western blots of the BN-PAGE-separated heart mitochondria of *Lrpprc*, *Tfam* and *Rnaseh1* knockouts (KO) and their corresponding controls (WT) presented in the main figures.
8. Interaction between ATP synthase and non-OXPHOS proteins or metabolic enzymes.
9. Distribution of interlinks of ATIF1 with ATP synthase proteins or metabolic enzymes.
10. Complexome profiling of wild-type mouse heart mitochondria shows the co-migration of ATPA, ATPB, MDHM and CISY.

#### Supplementary Figures

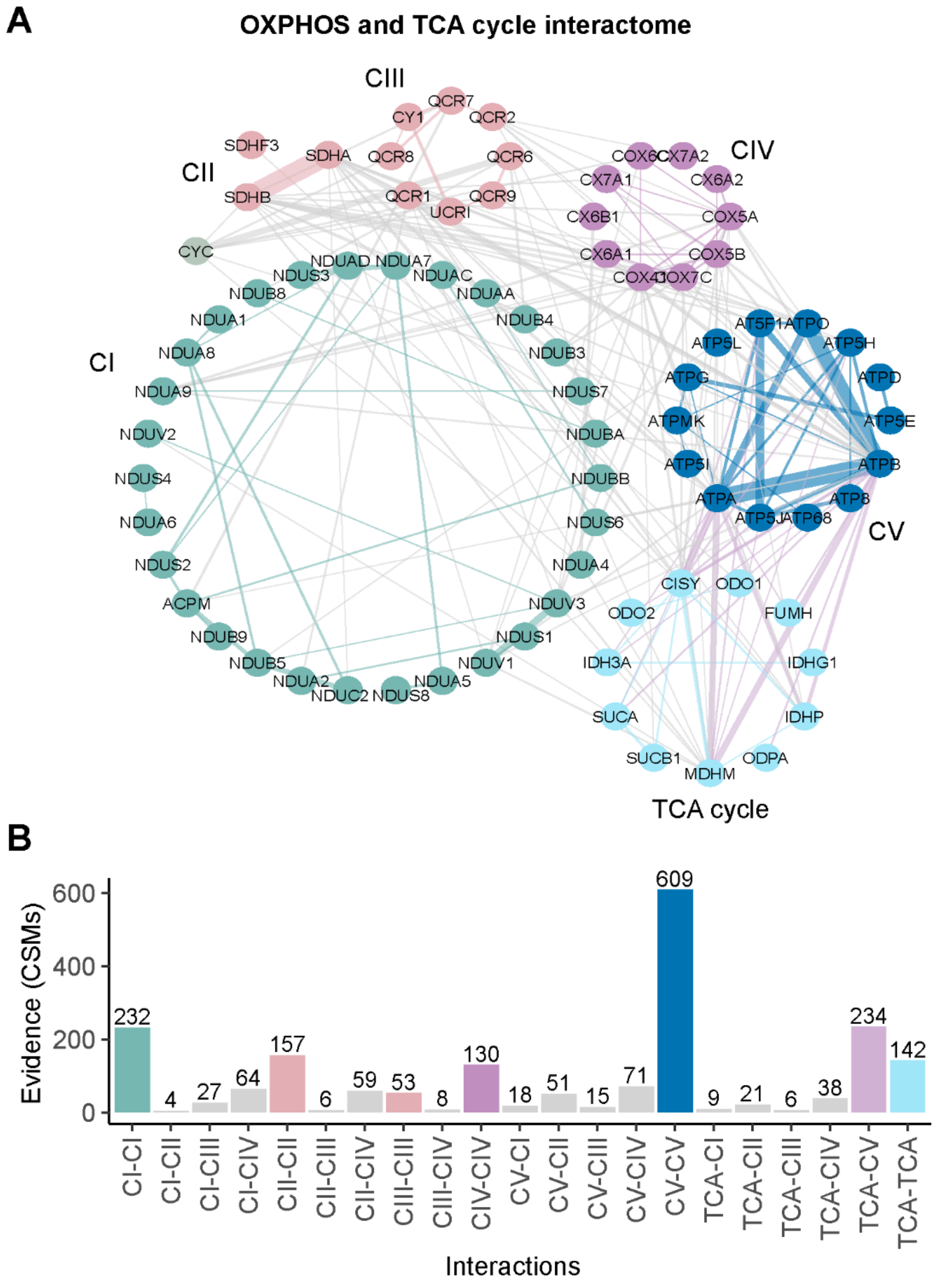


**Figure S1**. **XL-MS provides evidence for extensive interactions within and between the OXPHOS and TCA cycle proteins in mouse heart mitochondria.** (Based on data from Milenkovic *et al*, 2023). (A) Overview of protein interactions within the OXPHOS complexes and the TCA cycle, the names are abbreviated as follows: complex I (CI), complex II (CII), complex III (CIII), complex IV (CIV), ATP synthase (CV) and TCA cycle (TCA). (B) Bar plot of crosslink spectral matches (CSMs) detected between and within complexes across n=3 biological replicates.

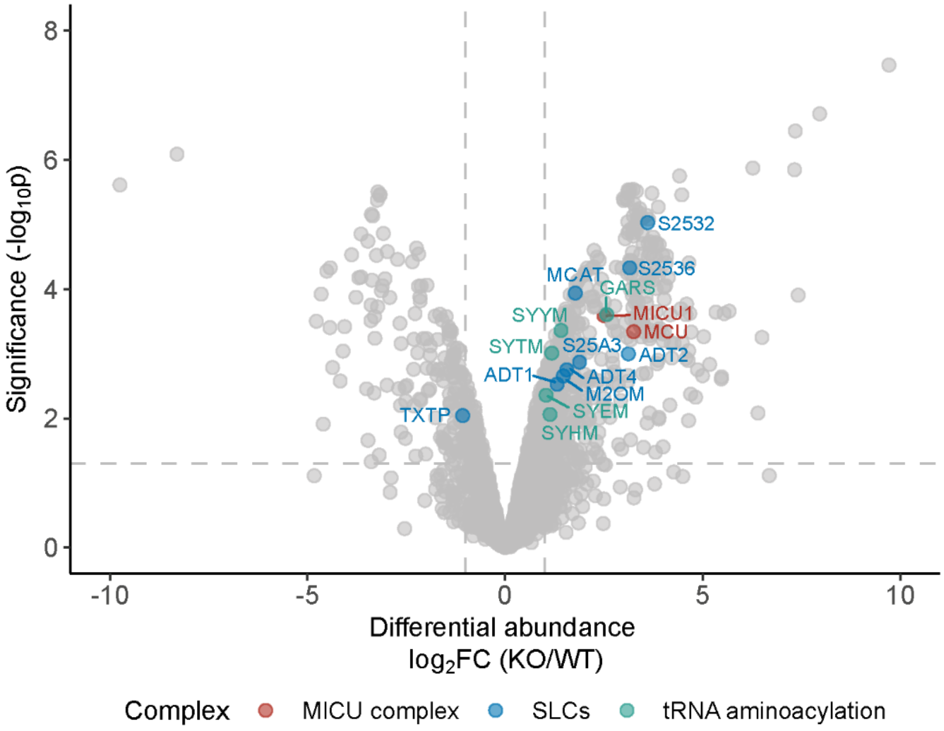


**Figure S2. Volcano plot highlighting previously undetected dysregulated mitochondrial proteins in heart mitochondria of *Lrpprc* knockout mice.** Differential expression analysis of protein abundances as detected in heart mitochondria of the *Lrpprc* knockout (KO) compared to the wild-type (WT) mice. The y-axis represents the-$\log_{10}$ p-value and the x-axis the $\log_2$ fold change (FC). The proteins from the mitochondrial calcium uptake (MICU) complex, mitochondrial solute carriers (SLCs) and tRNA aminoacylation-related enzymes are highlighted. Thresholds are an FDR < 5% and $|\log_2 FC| > 1$. Number of biological replicates, n=3.

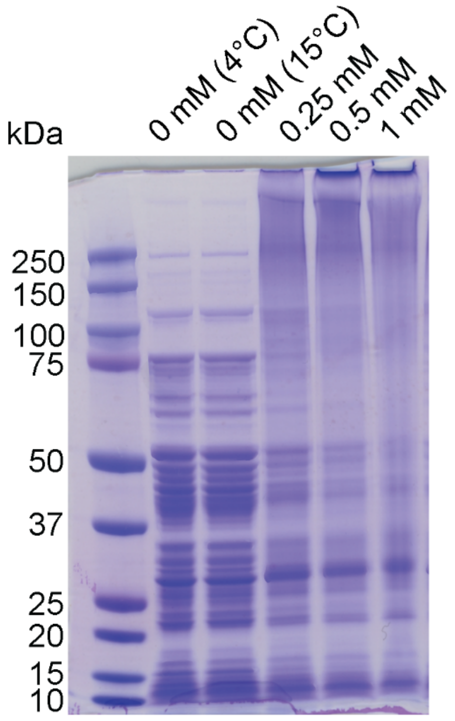


**Figure S3. SDS-PAGE used for the optimization of crosslinker concentration for the XL-MS experiments on mouse heart mitochondria.** Different concentrations (0.25, 0.5 and 1mM) were tested for the crosslinking reaction with DSSO. 0.5 mM was selected as the optimal concentration. The control reaction was tested both at 4ºC and at 15ºC to account for possible changes due to incubation temperature.

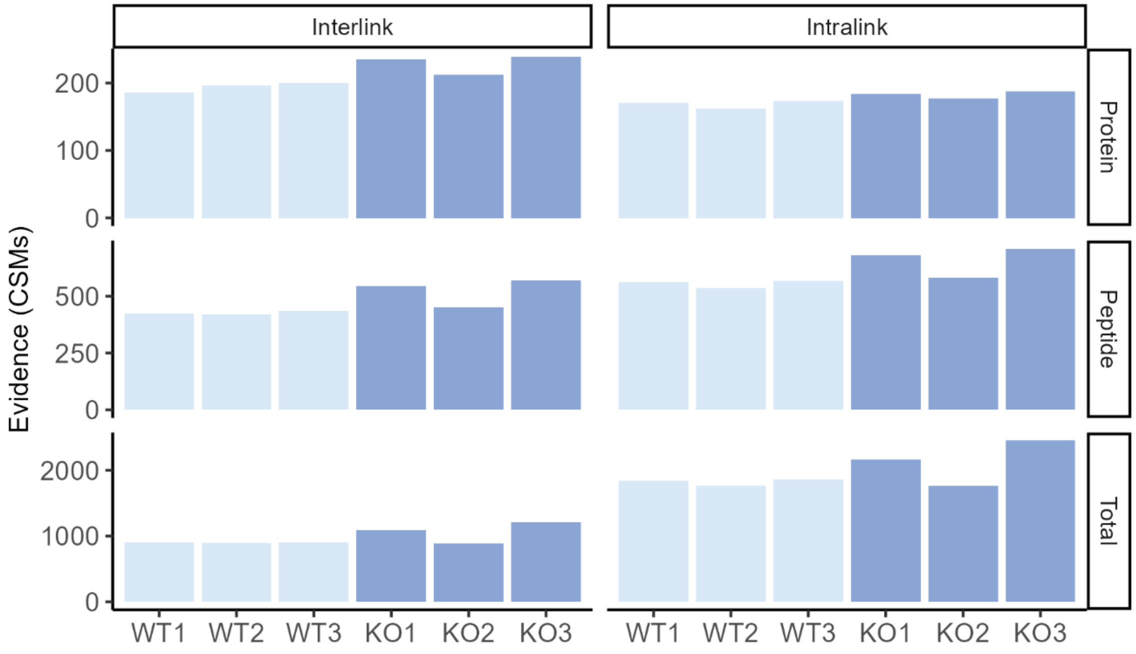


**Figure S4. Overview of CSMs detected per replicate in the XL-MS experiments**. Crosslink spectral matches (CSMs) identified per unique protein pair, peptide pair, and total. The values show good reproducibility across replicates.

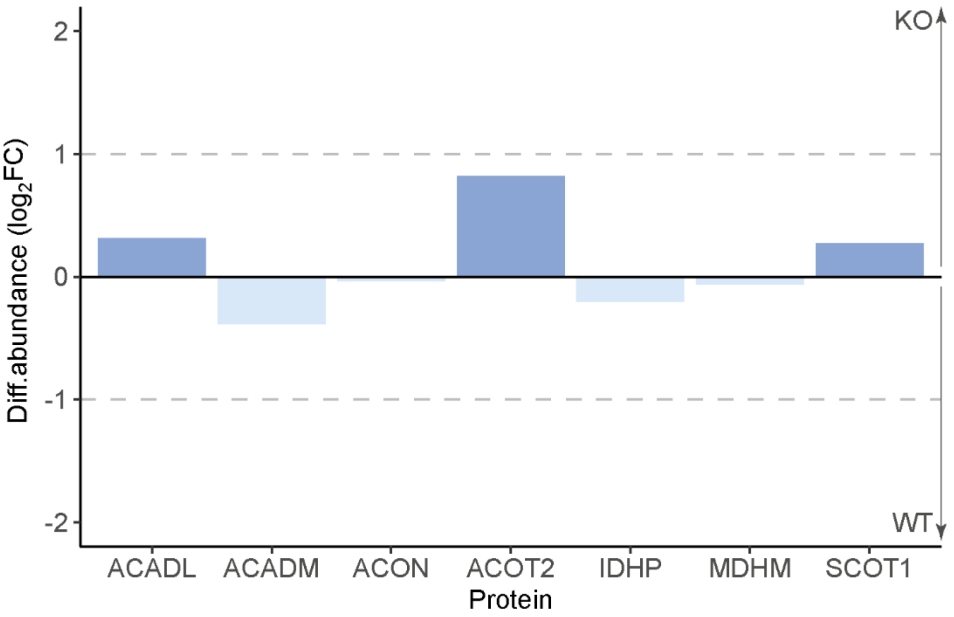


**Figure S5. Log$_2$ fold change (FC) of mitochondrial metabolism proteins up-regulated in the interactions with ATP synthase in the *Lrpprc* knockout hearts**. The fold change represents the difference between the protein abundances in the *Lrpprc* knockout (KO) and the wild-type (WT). All proteins had FDR > 5% and |Log$_2$FC| > 1 and were therefore considered not to be significantly changed. Number of biological replicates, n=3.

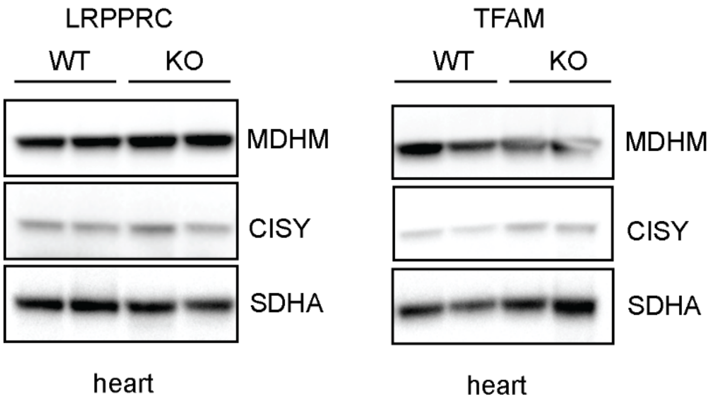


**Figure S6. Western blot analyses of SDS-PAGE-separated heart mitochondria from *Lrpprc* and *Tfam* knockout (KO) and their corresponding controls (WT).** No difference in abundance was observed in MDHM and CISY levels between the KO and WT mice. Minor differences might be caused by variability during sample loading.

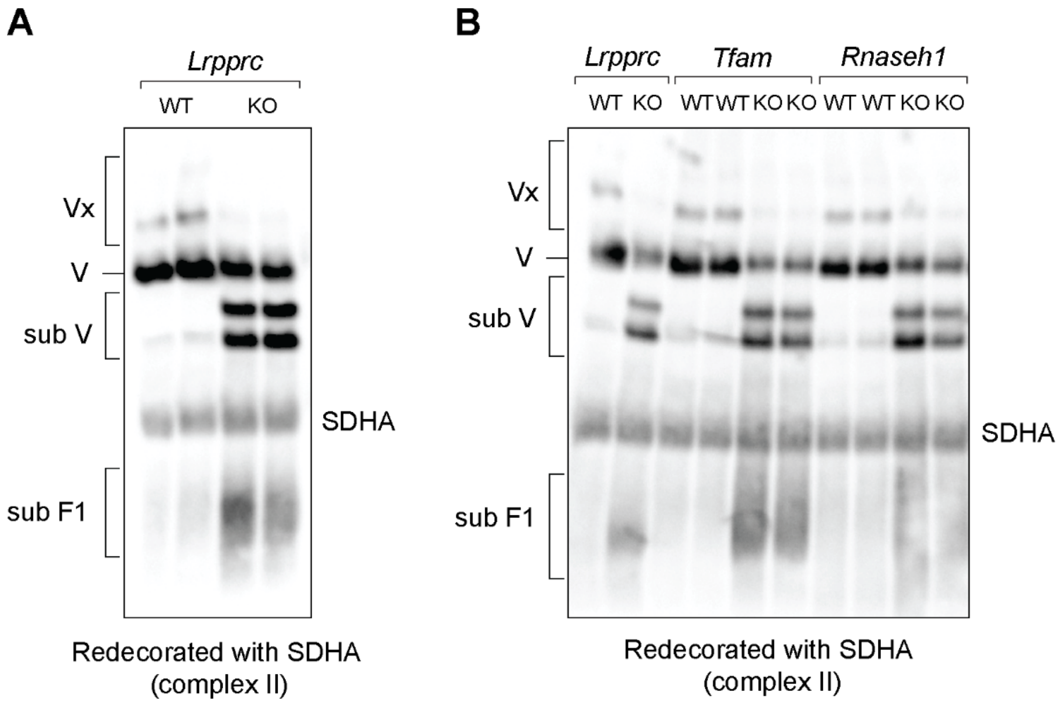


**Figure S7. Loading control for fluorescent western blots of BN-PAGE-separated heart mitochondria from *Lrpprc*, *Tfam* and *Rnaseh1* knockouts (KO) and their corresponding controls (WT) presented in the main figures.** Succinate dehydrogenase subunit A (SDHA, complex II) was used as the loading control for the fluorescent western blots shown in Figure 2F (panel A) and Figure 3B (panel B).

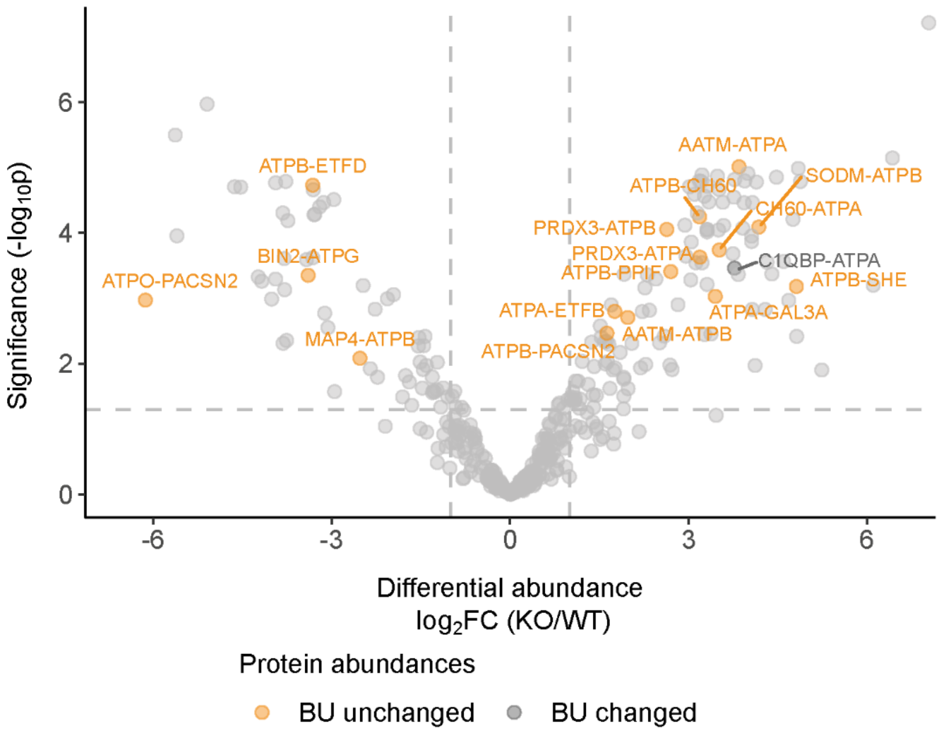


**Figure S8. Interaction between ATP synthase and non-OXPHOS proteins or metabolic enzymes.** Differential abundance analysis of the ATP synthase (CV) crosslinks with "other" proteins detected in the *Lrpprc* knockout (KO) compared to the wild-type (WT) heart mitochondria by XL-MS. The proteins annotated as "other" exclude all the OXPHOS structural components and metabolic enzymes, here interactions with ATIF1 were also excluded as they are presented in Figure 4A. Crosslinks involving proteins without significantly changed protein abundances according to the bottom-up (BU) proteomics analysis are highlighted in yellow; otherwise, they are colored in grey. The annotated crosslinks have an FDR < 5% and $|\log_2 FC| > 1$. Number of biological replicates, n=3.

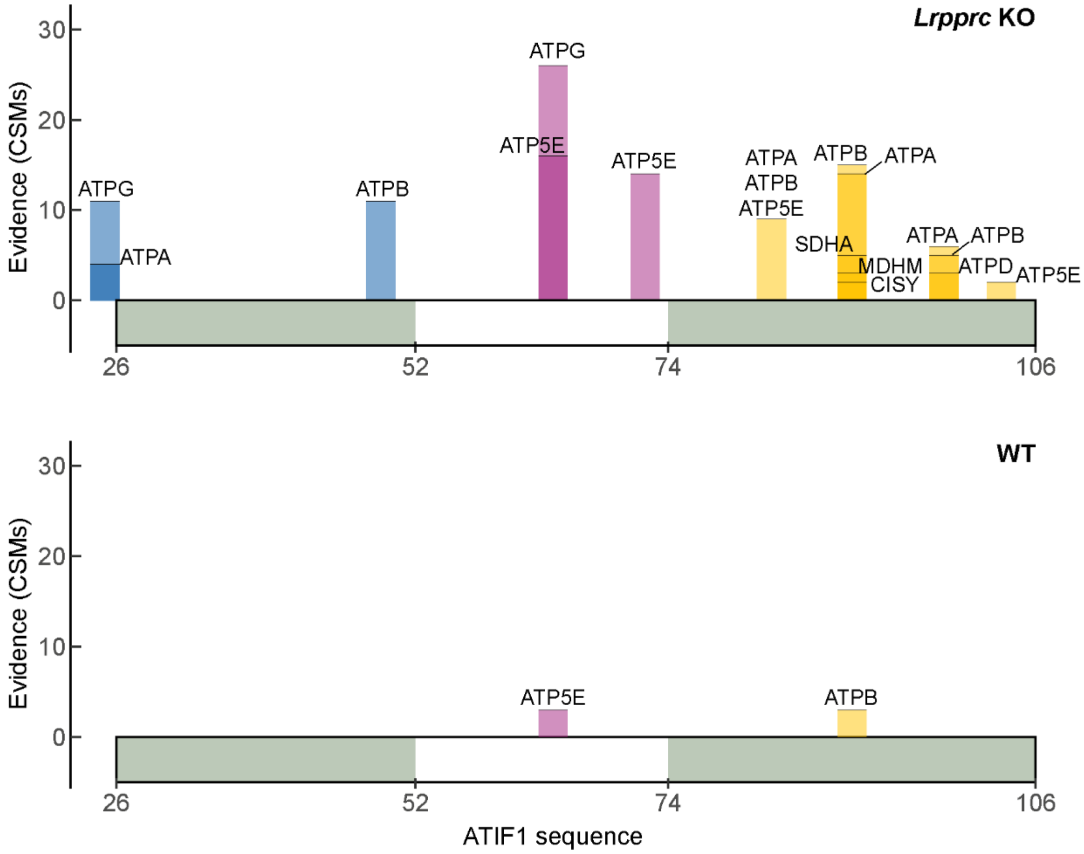


**Figure S9**. **Distribution of interlinks of ATIF1 with ATP synthase proteins or metabolic enzymes.** Interlinks mapped on the amino acid sequence of ATIF1 (without mitochondrial transit peptide), detected in the *Lrpprc* knockout (KO, top) and wild-type (WT, bottom) mice. For clarification, if different proteins are located at the same position on the bar, the number of detected crosslink spectral matches (CSMs) was identical. The CSMs represent the cumulative sum across n=3 biological replicates.

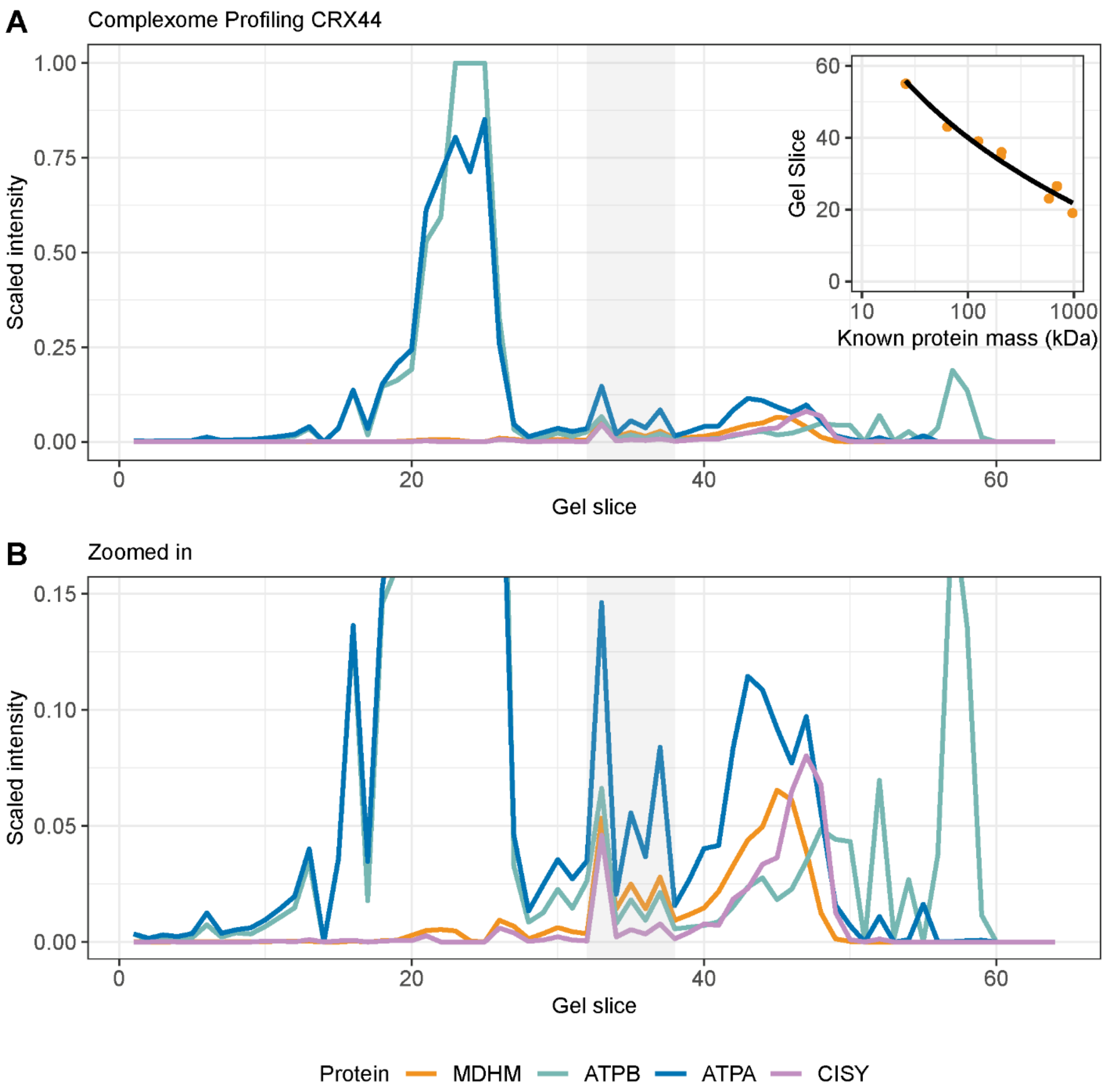


**Figure S10. Complexome profiling of wild-type mouse heart mitochondria shows the co-migration of ATPA, ATPB, MDHM and CISY.** The plot shows the protein intensities obtain by analysis of 64 gel slices by LC-MS/MS, from higher to lower molecular weight. Low Mw form of MDHM and CISY elute at gel slices > 40. The slices wherein these proteins co-migrate with ATPB and ATPA are highlighted in grey, the estimated molecular weight is in the range of approximately 275-140kDa. The molecular weight was estimated using the migration and masses of known complexes (see mass calibration graph). (B) shows the zoomed in view of (A). Data reanalyzed from *Yin et al,* 2024; available in the CEDAR database with accession number CRX44.
